# Microhabitat and ontogenetic variation of thermal tolerance in complex ectotherm life cycles

**DOI:** 10.1101/2022.12.14.520433

**Authors:** Jacinta D. Kong, Nicholas C. Wu

## Abstract

Thermal tolerances, such as critical temperatures, are important indices for understanding an organism’s vulnerability to changing environmental temperature. Differences in thermal tolerance over ontogeny may generate a ‘thermal bottleneck’ that sets the climate vulnerability for organisms with complex life cycles. However, a species’ microhabitat preference and life history can hinder our ability to assess climate change vulnerability in large-scale comparative studies as these reflect local adaptation. Here, we used phylogenetically informed, multi-level models with a global dataset of upper critical temperatures from 438 Anuran species that explicitly included microhabitat preferences and ontogenetic stage to examine variation in upper critical temperatures. We found ontogenetic trends in thermal tolerance were similar across microhabitat preferences. We then used microclimate-driven models of the heat and water budget of a thermoregulating frog to show the contribution of behavioural thermoregulation towards mitigating exposure to thermal stress. Our results suggested thermal bottlenecks are not strongly present in Anurans but instead implied strong developmental or genetic conservatism of thermal tolerance within families and ecotypes, and that behavioural thermoregulation has a strong potential for buffering frogs against extreme temperatures.

## Introduction

Thermal tolerance metrics, such as critical temperatures, have endured in ecological studies despite their criticisms because they are simple to conceptualise (e.g. impediment to movement), are generalisable across taxa, and can be readily assayed in the laboratory (Sinclair et al. 2016). These metrics facilitate our ability to link environmental temperature to organismal responses (Bates and Morley 2020), synthesise general trends in physiological diversity from local to macro scales (Chown and Gaston 2016), and index vulnerability or sensitivity to climate change via metrics such as thermal safety margins (Deutsch et al. 2008). For example, differences in upper critical temperature (CT_max_) across life stages may indicate a thermal bottleneck at the most temperature-sensitive life stage (Pandori and Sorte 2019, Dahlke et al. 2020). For organisms with a complex life cycle such as frogs, juvenile or larval stages in aquatic environments are generally expected to have lower thermal tolerance and thus be more vulnerable to climate change than adult stages (Lenoir et al. 2017).

Identifying a thermal bottleneck represents a significant amount of work that is limited to commonly studied species (e.g., Ruthsatz et al. (2022)). Thermal bottlenecks can also be identified using a macrophysiological approach which has described stage-specific differences in thermal sensitivities and tolerances in marine invertebrates (Truebano et al. 2018, Pandori and Sorte 2019, Collin et al. 2021), fish (Moyano et al. 2017, Dahlke et al. 2020), insects (Klockmann et al. 2017, Kingsolver and Buckley 2020) and reptiles (Levy et al. 2015). Among species comparisons in frogs are often limited to a single geographic region and are often limited to either adults or larva even if both are collected from the same region (Pintanel et al. 2019, Carilo Filho et al. 2021, Cheng et al. 2022, Pintanel et al. 2022, Ruthsatz et al. 2022). Thus, whether thermal bottlenecks occur and how consistent they are across Anuran species is not well understood. A macrophysiological analyses of thermal tolerance throughout ontogeny will vastly improve our understanding of the presence and generality of thermal bottlenecks across an environmentally sensitive taxonomic group.

Comparative macrophysiological analyses present two key challenges towards assessing and interpreting thermal tolerances. Microhabitat preferences and among-study variation may generate confounding variation in thermal tolerances that may mask or accentuate the presence of a thermal bottleneck. The key challenge to interpret thermal tolerance over the life cycle is to disentangle confounding variation deriving from microhabitat selection from inherent developmental or physiological drivers. A diversity of microhabitats may mask broader adaptive covariation between thermal tolerance and microclimate selection to explain, in part, an observed lack of variation in CT_max_ at a global scale or drive strong associations with thermal tolerance (González-del-Pliego et al. 2020, Pincebourde and Woods 2020). However, distinguishing between these patterns requires corresponding microclimate information for each ontogenetic stage that is not always collected for thermal tolerance studies or are available, but see Pintanel et al. (2022). If differences in thermal environments are consistent within broad microhabitat preferences in Anurans, then microhabitat preferences may be a reliable proxy for local environmental selection on thermal tolerance. Among species, frogs show strong microhabitat preferences that are associated with phenotypic syndromes encompassing behaviour, morphology and physiology and can broadly be categorised into ecological types (“ecotype”) (Moen and Wiens 2017, Moen et al. 2022). In turn, the animal’s ecotype may mediate the confounding interaction between life stages and their respective microenvironment (Pincebourde and Casas 2015, Carilo Filho et al. 2021).

Thermal safety margins (TSM) link upper critical temperatures to environmental temperature to index the vulnerability of ectotherms to temperature extremes (Deutsch et al. 2008). Most macrophysiological assessments of thermal safety margins use global datasets of climate and mean critical temperatures for species but the resolution of climatic data can have large impact on estimated TSM that leads to an underestimation of TSM (Garcia et al. 2019, Pinsky et al. 2019). Behavioural thermoregulation is important in this context as ectotherms that are able to behaviourally thermoregulate may be able to mitigate the effects of warming (Sunday et al. 2014). Thus, assessments of climate vulnerability using TSM should consider both the resolution of climate data and the innate capacity of species to respond to temperature variation.

Here, we tested whether ecotypes are reliable proxies of environmental adaptation to resolve ontogeny by environment interaction of CT_max_ and identify thermal bottlenecks. We analysed the CT_max_ of 438 Anuran species to 1) identify whether Anurans have a thermal bottleneck in their life cycle and their potential evolutionary drivers and ecological consequences at a macrophysiological scale, and 2) evaluate whether proposed solutions to two challenges of comparative analyses of ectotherm thermal tolerance improved our ability to identify thermal bottlenecks. We acknowledge there is a diversity of definitions and uses of thermal tolerance that would influence their interpretation. In this study, we only consider CT_max_ as the end point of a temperature ramping assay with a constant heating rate, or dynamic ramping protocol, and we define heating tolerance as the temperature range between starting temperature and CT_max_ during the ramping assay.

## Methods

### Data compilation

We started with a published dataset of upper critical temperatures of Anurans complied from the literature (1962 – 2015) (Rohr et al. 2018). We supplemented this data with studies from 2016 to 2022 using Web of Science (accessed 25th February) and the keywords amphibian* AND (“critical thermal max*” OR CT_max_ OR “thermal toleran*” OR “critical temp*” OR “upper temp*”)”. The terms returned 90 studies that were screened for inclusion by one person (J.D.K.). Studies that measured CT_max_ using ramping assays were retained. Studies that estimated CT_max_ from fitted thermal performance curves or compiled values from literature were excluded. Only control animals were included from studies that used multiple stressors (e.g. dehydration or infection). The screening process yielded 36 papers that provided extractable data. Provided data were used over published values where possible. Populations were considered independent observations. Multiple endpoints of critical temperatures on an individual were recorded but only one CT_max_ was used per individual (loss of righting response where possible). We added studies that did not return from keywords from forward searching. We distinguished between temperatures animals were held at prior to experimentation (acclimation temperature) and the starting temperature of the ramping assay. We assume that observed CT_max_ reflected their acclimated CT_max_ as acclimation in frogs is reported on scales of hours (Brattstrom 1970).

We obtained ecological types (hereby ecotype), from all Anuran species listed on the International Union for Conservation of Nature’s Red List of Threatened Species (IUCN 2022). The data were reported in (Wu et al. 2024). Ecotypes were classified from ecological descriptions from IUCN based on adult behaviour and microhabitat preferences outside the breeding season (Moen and Wiens 2017). We grouped species by ecotype even if the larval stage may have differing ecological types (e.g. lotic vs lentic systems) because the ecotype diversity of the larvae stage is broader than adults and is less described in the detail from the literature. The resulting dataset represented 32 families spanning the globe (Fig. S1). Thermal tolerance records were first matched to the ecotype dataset by binomial names used in the original source, then unmatched records were joined by IUCN names based on synonyms accepted by IUCN or Catalogue of Life. CT_max_ records that could not be matched with ecotype (56 observations) were excluded.

### Relationship between CT_max_ and heating tolerance with ecotype and ontogeny

We constructed a phylogenetically corrected multi-level Bayesian model to investigate the relationship between CT_max_ or heating tolerance across ontogenetic stage and ecotype (see Supplementary Information for detail). We included starting temperature of the assay as a co-variate. Ontogenetic stages 1-5 correspond to larva, 6 to juveniles, and 7 to adults (Ruthsatz et al. 2022). We used an amphibian phylogeny dataset comprising of 7,238 species to account for phylogenetic relatedness (Jetz and Pyron 2018). We pruned the phylogenetic tree to matched binomial names in the thermal tolerance data (438 species, Fig. 2). Species names based on Jetz and Pyron (2018) were used in analysis to account for duplicate names matching with the IUCN. Records that were not in the phylogenetic tree (n = 1) were excluded. We converted the pruned tree to a phylogenetic relatedness correlation matrix for subsequent analysis. For the CT_max_ model, we used a quadratic equation to model change in CT_max_ over ontogeny and the quadratic equation provided a better fit than the linear model (Table S1). All model outputs (Tables S2 – S5) and posterior predictive checks (Figs. S4 & S6) are presented in the Supplementary Information. All model parameters are reported as mean Bayesian posterior estimates ± 95% credible intervals (95% CI), which is equivalent to 95% confidence intervals for frequentist statistics. Summary statistics are presented as mean ± standard deviation.

### Thermal safety margins at macro- and micro-climate scales

To calculate thermal safety margins we extracted mean annual temperature and maximum temperature of the warmest month from WorldClim 2.1 (resolution: 0.5 min of a degree) for each observation (Fick and Hijmans 2017). TSM was calculated as upper critical temperature (CTmax) - maximum temperature of the warmest month. Observations that could not be queried on WorldClim were removed from analysis. To illustrate the effect of spatial resolution on estimates of TSM for representative ecotypes, we chose frogs representing three ecotypes (Fig. 1): fossorial (*Neobatrachus pictus*), arboreal (*Litoria peronii*) and ground-dwelling (*Crinia signifera*). These species are sympatric in north-eastern New South Wales, Australia and were measured in the same study using the same conditions (5°C acclimation temperature and 1°C min^-1^ heating rate) (Brattstrom 1970). Because of the additional challenges of modelling microclimates involving water and the potentially large temperature difference between air and water, we were not able to conduct simulations for aquatic, stream-dwelling and semi-aquatic ecotypes We used NicheMapR to simulate representative microclimates for a fossorial (90% shaded 0.1 m soil temperature), arboreal (60% shaded 1m air temperature) and ground-dwelling species (0% shaded 0.01m air temperature) (Kearney and Porter 2017). These microclimates informed a biophysical energy and mass balance model of a hypothetical non-thermoregulating 5g frog (approximating a leopard frog) that we used to estimate operative temperature (maximum body temperature) for each species (Kearney and Porter 2019, Enriquez-Urzelai et al. 2020, Wu et al. 2024). To demonstrate the role of behavioural thermoregulation for estimating TSM, we ran the biophysical model twice for the ground-dwelling frog allowing for or restricting microhabitat selection by shuttling between shaded and unshaded microhabitats. For each ecotype, we extracted maximum temperature at the locality of the individual and maximum body temperature, representing operative temperature, from the year-worth of output (Fig. 1).

**Figure 1.**
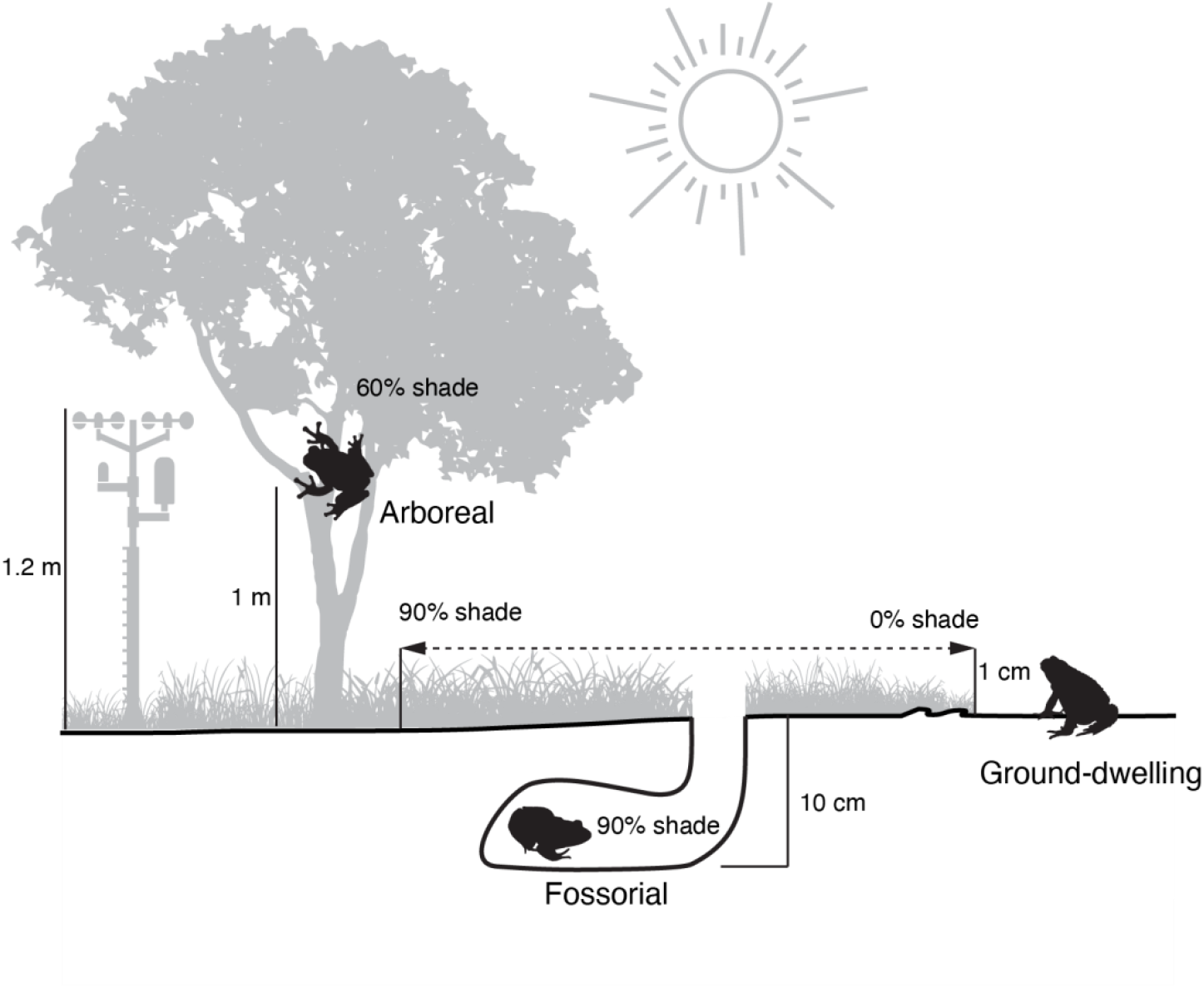
Schematic diagram of ecotype microclimate and key variables used to calculate representative microclimate. Arboreal frogs were modelled at 1m 60% shaded air temperature, fossorial frogs at 10cm soil depth in 90% shaded soil, and ground-dwelling frogs were modelled at 1cm 0% shaded (open) air temperature. Thermoregulating ground-dwelling frogs could move to 90% shaded 1cm air temperatures, indicated by the arrows.

**Figure 2.**
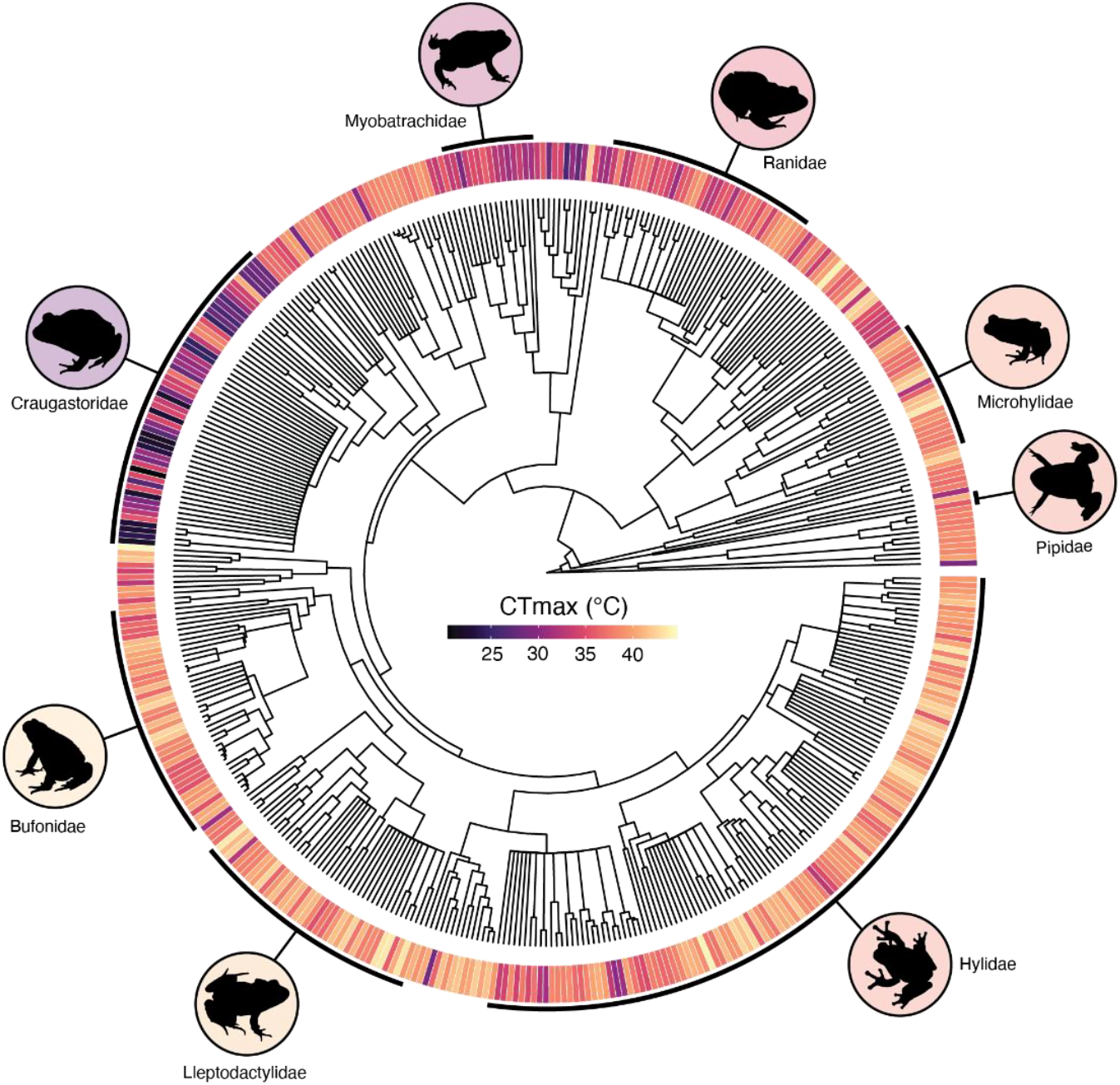
Phylogenetic patterns of CT_max_ across 438 Anurans based on Jetz & Pyron (2018). Colours represents species mean CT_max_. Example families are indicated by outer black lines. Representative silhouettes obtained from phylopic.org and background colours within the circles represent mean CT_max_ at the Family level.

## Results

### CT_max_

CT_max_ ranged between 20.4 °C and 46.1 °C (mean 37.1 ± 4.1 °C) across all ecotypes. On average, fossorial frogs have the highest mean CT_max_ (40.1 ± 3.1 °C) and semi-aquatic frogs have the lowest CT_max_ (35.4 ± 3.54 °C). Larval CT_max_ (Stage 1-5) was on average 3.5 °C higher than adult (Stage 7) and juvenile (Stage 6) CT_max_ combined; the mean difference ranged from 1.5 °C for semi-aquatic frogs and 5.2 °C for stream-dwelling frogs. However, there are fewer data on CT_max_ for early larval (Stage 1, n = 41) and juvenile (Stage 6, n = 33) stages than later larval stages (Stage 3, n = 207) and adults (Stage 7, n = 733).

Variation in CT_max_ was strongly influenced by ontogeny where CT_max_ generally decreased across ontogeny (Table S2; Fig. 3). CT_max_ did not differ substantially between ecotypes, but on average, stream-dwelling frogs have lower CT_max_ (−0.74 [-2.63: 1.11]), while fossorial frogs generally had higher CT_max_ (1.96 [-0.06:4.05]). There was a strong phylogenetic effect on CT_max_ (λ = 0.85 [0.75:0.92]). Starting temperature also had a strong effect on CT_max_, where the higher the starting temperature, the higher the CT_max_ (0.16 [0.14:0.18]).

**Figure 3.**
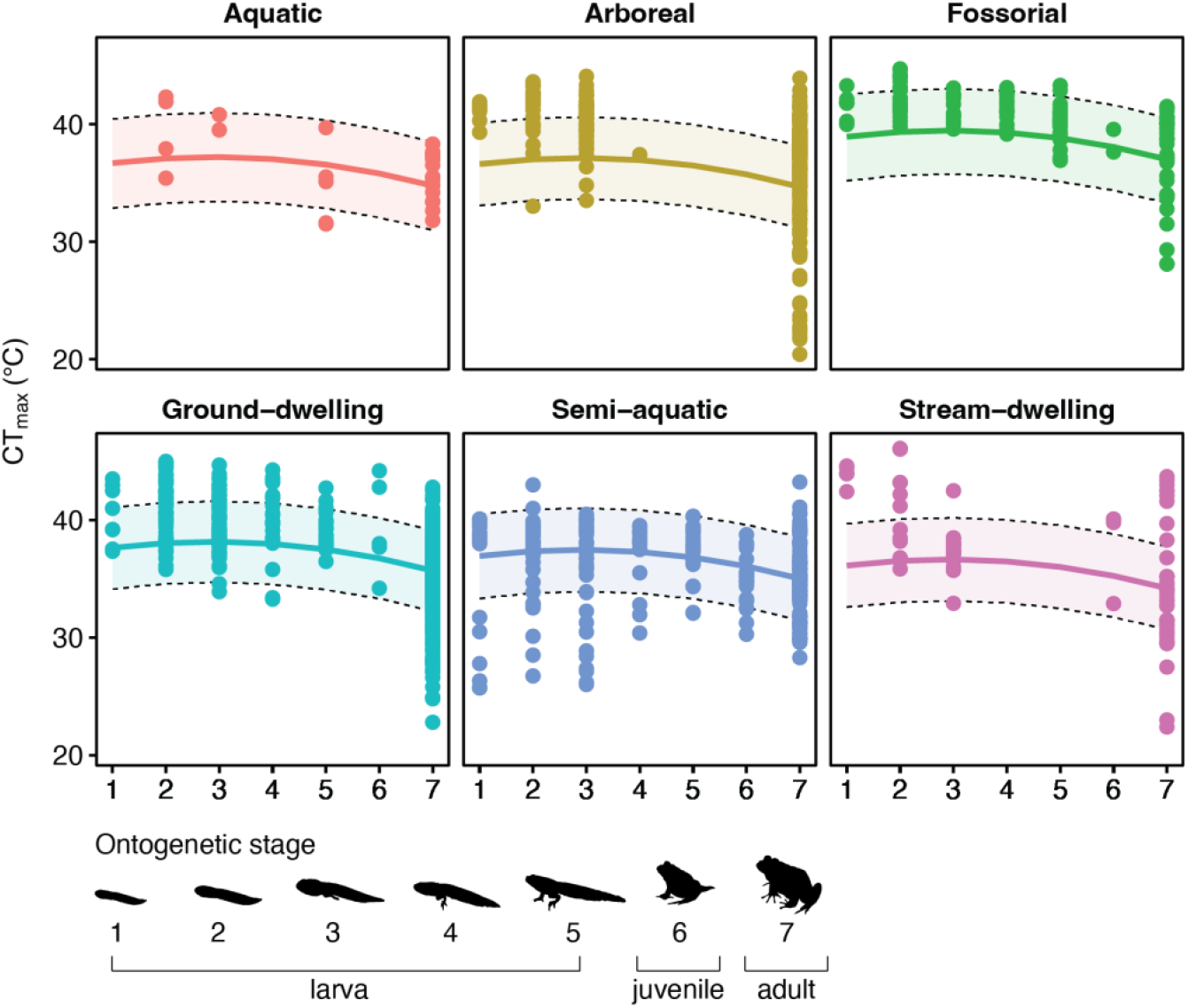
CT_max_ (^°^C) decreases over ontogeny (stage 1–7) across all Anuran ecotypes (438 species). Quadratic regression line and shaded area represents ecotype-specific model estimate ± 95% CI.

### Heating tolerance

Heating tolerances ranged between 2 °C and 36.5 °C (16.7 ± 6.34 °C) across all ecotypes. Mean heating tolerance varied between ecotypes with the highest mean in fossorial frogs (18.6 ± 6.1 °C) and lowest in stream-dwelling frogs (16.0 ± 6.2 °C). Heating tolerance was also influenced by ontogenetic stage (−0.42 [-0.49:-0.35); Fig. 4), with marginal influence by ecotypes (Table S4). There was a strong phylogenetic effect on heating tolerance (λ = 0.86 [0.76:0.92]) and an effect of starting temperature where heating tolerance was lower at higher starting temperature (−0.83 [-0.85:-0.82]).

**Figure 4.**
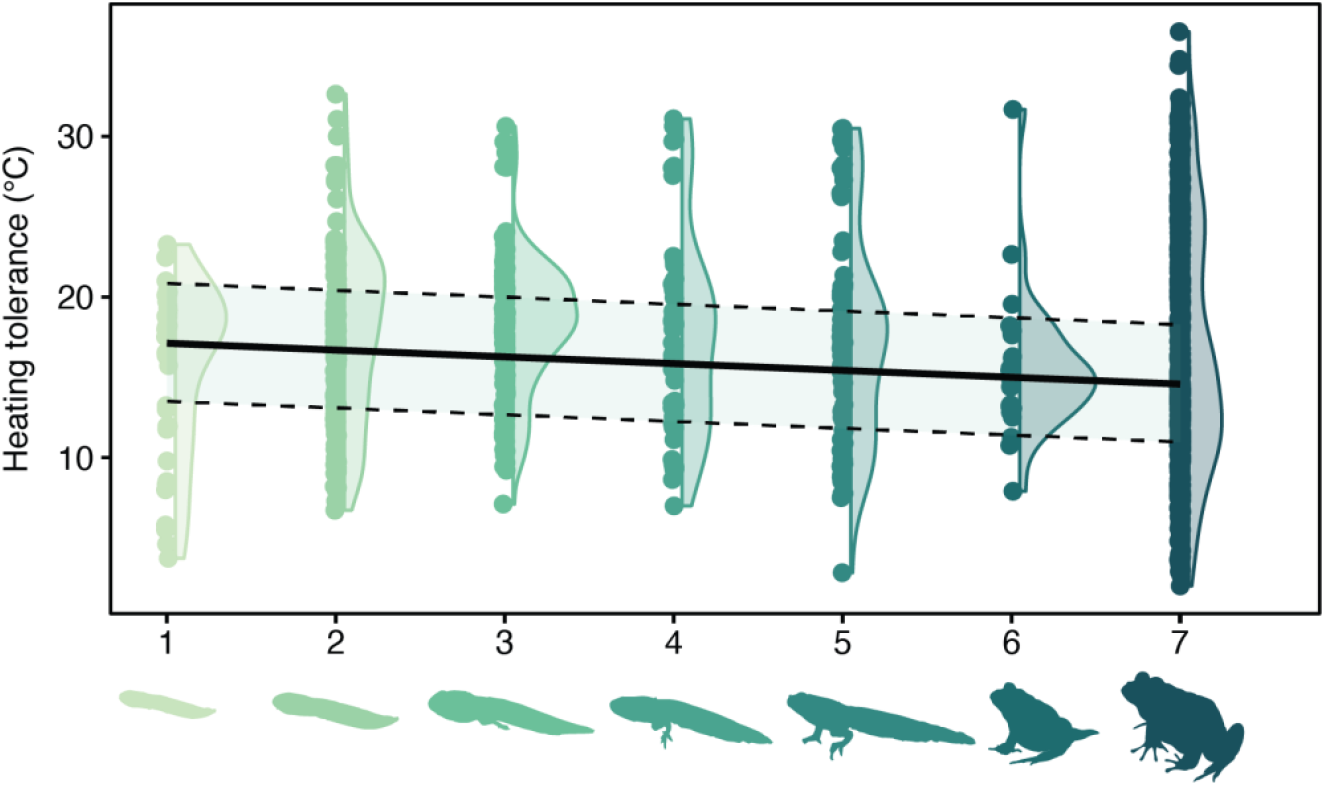
Heating tolerance (°C) decreases across ontogeny (stage 1–7, colours). Regression line and shaded area represents overall model estimate + 95% CI. Heating tolerance is CT_max_ – starting temperature of the ramping assay (438 species).

### Thermal safety margin

Estimates of thermal safety margin vary with the spatial resolution of the environmental temperature used in calculations, microhabitat characteristics, and the presence of behavioural thermoregulation (Table 1, Fig. 5). Using macroclimate underestimates TSM for fossorial and ground-dwelling frogs because the microclimate of these ecotypes are not well captured by weather stations. Higher TSM using microclimate than macroclimate for a fossorial species are generated by a lower soil temperature than air temperature. There is little difference between using operative temperatures and microclimate because fossorial frogs do not leave their burrow. TSM are similar using all spatial resolutions for arboreal frogs because arboreal microhabitats represented by 1m shaded environments (60% canopy cover) are similar to weather stations (e.g. 1.2m 90% canopy cover). A ground-dwelling frog in an open, sunny habitat will encounter maximum air temperatures and exhibit maximum body temperatures exceeding their upper critical temperature (negative TSM) at the relevant microclimatic scale of the individual (1cm above ground in our simulations). When allowed to select cooler microhabitats, predicted body temperatures of simulated ground-dwelling frogs never exceed CTmax (positive TSM).

**Table 1.**
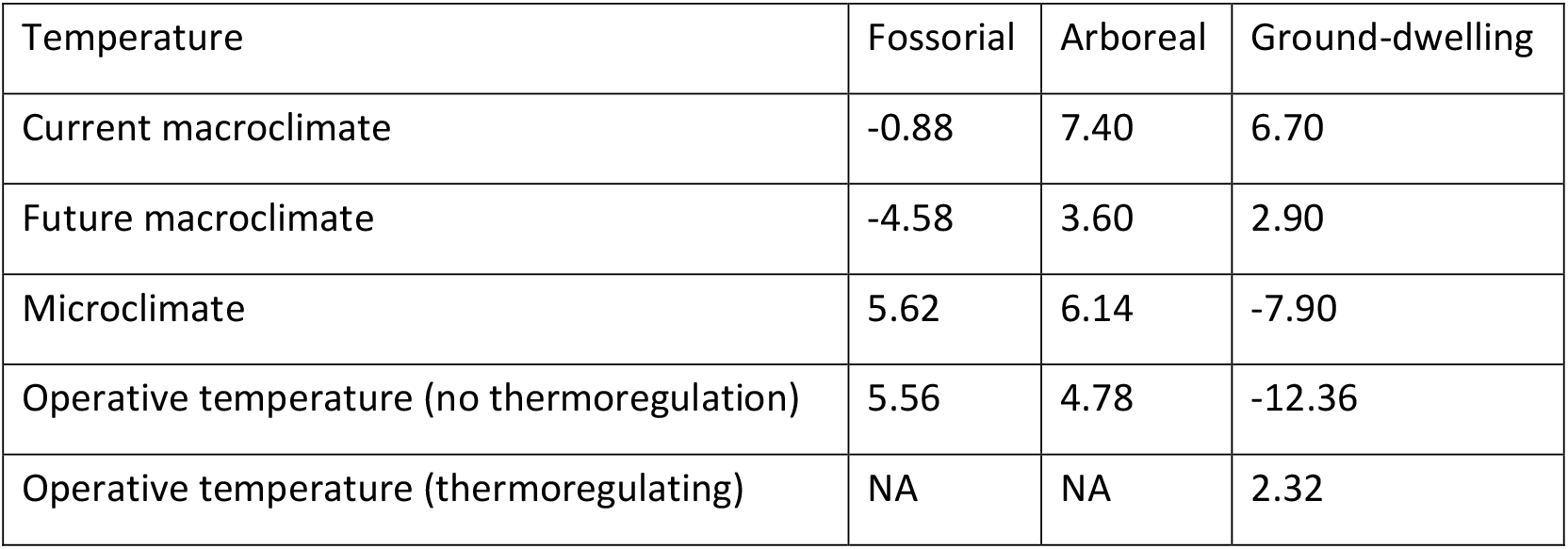
Thermal Safety Margin (TSM; CTmax – maximum temperature; C) calculated using maximum temperatures of varying spatial resolution for sympatric species representing three ecotypes: fossorial (Neobatrachus pictus), arboreal (Litoria peronii) and ground-dwelling (Crinia signifera) species. Macroclimate: Maximum temperature of the warmest month (WorldClim, BIO5). Microclimate: Maximum temperature from downscaled climate to representative microhabitat conditions of the ecotype (NicheMapR). Operative temperature: maximum body temperature for a simulated 5g frog in their preferred microhabitat.

**Figure 5.**
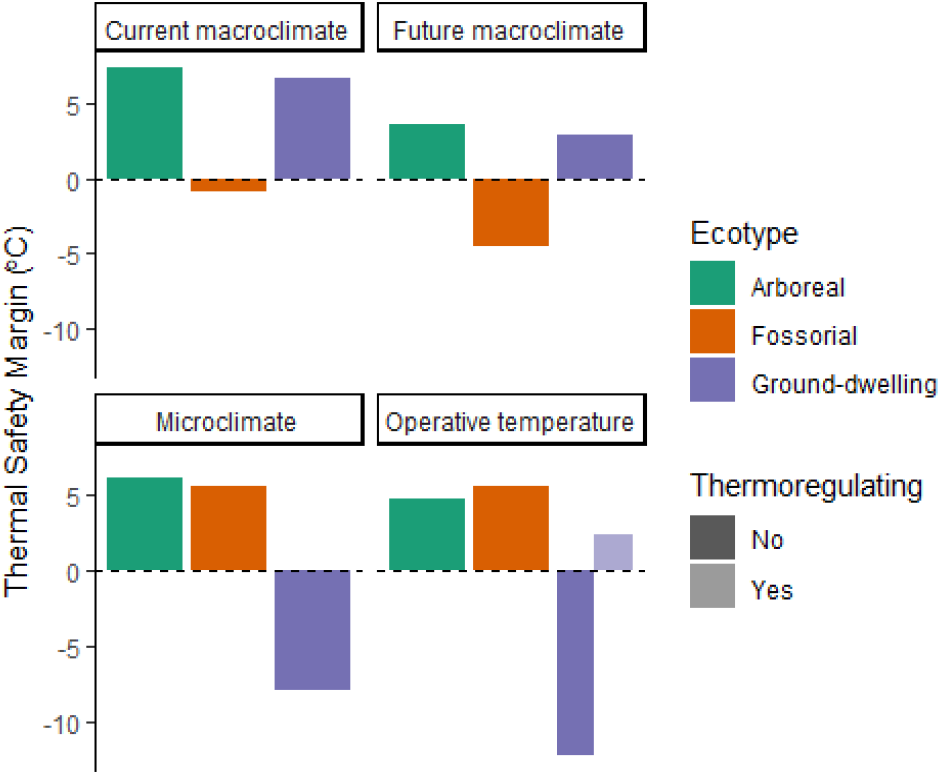
Thermal Safety Margin (TSM; CTmax – maximum temperature; C) calculated using maximum temperatures of varying spatial resolution for sympatric species representing three ecotypes: fossorial (Neobatrachus pictus), arboreal (Litoria peronii) and ground-dwelling (Crinia signifera) species. Macroclimate: Maximum temperature of the warmest month (WorldClim, BIO5). Microclimate: Maximum temperature from downscaled climate to representative microhabitat conditions of the ecotype (NicheMapR). Operative temperature: maximum body temperature for a simulated 5g frog in their preferred microhabitat.

## Discussion

Amphibians are an excellent system in which to investigate ontogenetic variation in thermal tolerance because they have a biphasic life cycle with life stages clearly delineated by metamorphosis. The delineation between life stages set by metamorphosis is key as metamorphosis may compartmentalise the life cycle and allow for a potential thermal bottleneck at one stage with minimal detrimental effects of temperature across a life cycle (Enriquez-Urzelai et al. 2019, Lowe et al. 2021).

We found striking differences between adults and larva across ecotypes in the opposite direction than expected if larva represented a thermal bottleneck. A similar general negative relationship between thermal tolerance over ontogeny is widely reported for frogs (Cupp 1980, Enriquez-Urzelai et al. 2019, Ruthsatz et al. 2022). Although lower thermal tolerances may be expected for aquatic larva, a higher thermal tolerance may be advantageous for small bodied larva that have reduced capacity to escape temperature extremes (Lenoir et al. 2017). Similarly, a wider selection of microhabitats in terrestrial environments may allow adults to compensate for a lower thermal tolerance through behavioural site selection than aquatic larva which grow in a thermally homogeneous environment (Enriquez-Urzelai et al. 2020).

We found ecotype mediates the magnitude of ontogenetic differences in thermal tolerances but is not an accurate proxy for environmental selection or constraint. The discrete nature of complex life cycles with metamorphosis and the need to survive thermal stress at one stage until the transition to the next stage may place high importance on the interaction between traits and the environment where a thermal bottleneck is a selective disadvantage (Lowe et al. 2021). Microhabitat preferences has been demonstrated to strongly correlate with thermal biology in reptiles (Greer 1980, Michelangeli et al. 2018), bees (Wilson et al. 2020), frogs (Duarte et al. 2012, Carilo Filho et al. 2021), and ants (Bujan et al. 2020, Leahy et al. 2022). We found strong phylogenetic conservatism in CT_max_ which likely reflects how ecotypes strongly correlate within Family (Moen and Wiens 2017). For example, Hylidae have a high proportion of arboreal species (Vidal-García and Keogh 2015). The use of ecotype as a proxy is complicated by variation in developmental characteristics that may not be consistent within Family or ecotypes to make generalisations. For example, direct-developing larva (not available in our dataset) may occupy more similar microhabitats to adults than species with an aquatic larva and fully terrestrial adults as discussed with the limitations of larvae ecotypes. Egg thermal tolerance is rare in the Amphibian literature, but eggs may also occupy vastly different microhabitats to larva and adults. For instance, arboreal frogs may create foam nests in the canopy but the larva inhabit puddles on the forest floor (Haddad and Prado 2005). While it would be possible to classify larval ecotypes or egg microhabitats (Carilo Filho et al. 2021), this information is not readily available at the scale of this study, and we encourage further investigation into these modulators. Doing so may reveal whether diversity in developmental traits within ecotypes masks local scale signals of environmental adaption via interactions with ecotype. Nevertheless, we found fossorial species, on average, had higher thermal tolerance than other ecotypes.

The large overlap in thermal tolerance across ecotypes suggests developmental or genetic constraints have a greater influence on the evolution of thermal tolerances than environmental factors. Developmental or genetic constraints on climate niches or thermal traits are suggested for reptiles and amphibians (Pie et al. 2017, Bodensteiner et al. 2021, Carilo Filho et al. 2021). Several inferences support developmental or genetic constraints underlie consistent differences between larva and adult thermal tolerance. First, an absence for a bottleneck may indicate metamorphosis modulates thermal tolerances over ontogeny (Lowe et al. 2021). Minimal carry over effects of development temperature on CT_max_ are found between larva and adults (Enriquez-Urzelai et al. 2019). Physiological remodelling and morphological development during metamorphosis are hypothesised to trade-off with physiological processes and explain why CT_max_ decreases at metamorphic climax (Cupp 1980, Ruthsatz et al. 2022).

Second, if juveniles and adults are not drastically different in their physiology (i.e., biochemical processes are not affected by metamorphosis), then differences in thermal tolerance may indicate differences in energy allocation between life stages. An energetics interpretation depends on whether capacity to respond to a thermal challenge is considered part of maintenance metabolic costs and the flexibility of reallocating energy. Perhaps, the allocation of energy to reproduction at the adult stage trade-offs with the capacity to increase maintenance metabolic rates in response to heating, whereas larva without a reproductive metabolic cost have greater scope to rapidly respond to a thermal challenge at the cost of allocating energy to growth. This explanation is in line with how the temperature size rule emerges from differences between development and growth rates (Forster and Hirst 2012, Rubalcaba and Olalla-Tárraga 2020, Gahm et al. 2021).

A higher physiological capacity to respond to change (i.e., plasticity) in larva is a parsimonious explanation from an energetics perspective. In contrast with studies that show decreasing acclimation capacity with increasing size (Rohr et al. 2018, Burton et al. 2020), *Rana temporaria* larva have lower acclimation capacity than adults (Ruthsatz et al. 2022). Assuming the cost of having a given thermal tolerance is the same between life stages, low plasticity in thermal tolerance suggests larva continuously express physiological mechanisms that permit higher CT_max_, which is very costly (i.e., in maintenance metabolic cost) during a time when they should be allocating energy to growth. One explanation for low thermal plasticity is tadpoles have a narrow range of temperatures to select for behavioural thermoregulation and thus primarily rely on physiological thermoregulation via a higher CT_max_ to avoid heat stress (Casterlin and Reynolds 1978). As larva would have lower absolute metabolic rate based on size allometry alone (in general), whether or not they can offset an increased energetic cost with increased food intake or by other means can be explored.

Third, if physiological mechanisms are different between life stages, then the differences may reflect developmental constraints. Perhaps the physiological machinery during larval growth covaries with greater capacity to respond to an environmental challenge, or that larva have lower energetic costs for responding to thermal challenges, which allows them to continuously express high CTmax at minimal cost to growth, in contrast with adults. This explanation would resolve the paradox of the high metabolic costs of physiological thermoregulation.

A notable characteristic of the amphibian CT_max_ literature is that there is a relative consistency in methods to assay CT_max_ and the range of heating rates is narrow with a mode of 1 °C min^-1^ (Hutchison 1961). Most studies in our literature search (2016-2022) assayed CT_max_ using a ramping protocol rather than static protocols (zero studies described) or from fitted thermal performance curves (three studies). In contrast, there is a much larger diversity in heating rates reported for global ectotherm CT_max_ varying between 0.0007 and 4°C min^-1^ (mean 0.5 °C min^-1^) that reflects the tropical to polar origin of species (Morley et al. 2018). In studies on frogs, ontogenetic stage is confounded with heating rates as lower heating rates are typically used for larva than adults or juveniles (Fig. S3). This different may contribute towards nuisance heterogeneity in comparative study but the effects may be minimal compared to other taxa (Noble et al. 2022). In our dataset, CT_max_ endpoints are relatively consistent across studies and within ontogenetic stage, but ontogenetic stage was confounded by endpoint definition. For example, onset of spasms or heat rigour were more common for larva and loss of righting response is most common for adults. A model invoking a hypothesised mechanism could be relevant for investigating specific endpoints or definitions of upper critical temperatures.

Simulating body temperatures trades-off with making significant assumptions about the microclimate characteristics of each ecotype and their thermal biology. For example, arboreal frogs bask but we simulated they did not (Tracy et al. 2010). Using operative temperature is best suited to species with well characterised thermal biology to derive accurate TSM estimates and activity patterns at a local scale, for species whose body temperatures are expected to be significantly different to environmental temperature, or for large-scale studies that trade-off accuracy for scale. If the thermal biology of species is known, then a microclimate-driven biophysical models of ectotherm body temperatures can provide a fine-scale analysis of estimated activity times under warming scenarios and hypothetical capacities to mitigate potential heat stress (Anderson et al. 2022).

A size-dependency of CTmax may explain why heating tolerances are relatively conserved over ontogeny. Larger sizes may buffer adult frogs against higher heating rates via inertia effects, thus a lower CTmax for an adult is equivalent to higher CTmax in smaller larva when rate-time dynamics are considered. A lower size limit for thermal inertia in dry-skinned ectotherms is approximately 1kg, which is larger than most frogs. A similar threshold for frogs is unclear. While tadpoles are reported to be at thermal equilibrium with water temperature, a small basking arboreal frog (<10g) can elevate their body temperature above ambient temperature. Rates of heat exchange that influence body temperature changes in frogs (a wet-skinned ectotherm) depends on resistance to water loss, size, and the difference in thermal properties between water and air.

Microclimate and microhabitat preference strongly modulate estimates of climate vulnerability. We demonstrate that spatial resolution of temperature strongly influences estimated metrics of climate vulnerability. The magnitude of under- or over-prediction was highly dependent on the microhabitat preferences of species (Table 1). Specifically, canopy cover, which intercepts direct solar radiation, in the microclimate model set the ecological accuracy of the simulated microhabitat and may differ from the selected microhabitats in the field. In general, we suggest using simulated microclimates to estimate TSM than macroclimate, particularly for large scale studies for which field measurements of temperature are not feasible. Doing so relies on fewer assumptions than using a biophysical model of the heat balance of a generalised frog to estimate operative temperatures, requires one model instead of two for deriving environmental temperature to calculate TSM, and may be a reasonable proxy for body temperature for an ecotype. It may be reasonable to use soil temperature as a proxy for body temperature for fossorial frogs that do not frequently leave their burrow.

Our simplified simulations clearly illustrate the importance of behavioural thermoregulation (Bogert effect) and the availability of thermal refugia in the context of assessing thermal safety margins (Kearney 2013, Sunday et al. 2014). At face value, our results suggest for a given location, ground dwelling frogs are highly vulnerable to climate change, but the frog was not simulated to avoid temperatures above their activity thresholds. Several studies demonstrated microclimate is a better predictor of frog CTmax than microclimate (Gutiérrez-Pesquera et al. 2016, Katzenberger et al. 2018, Pintanel et al. 2019), and that variation in tadpole CTmax is explained by local scale water temperatures measured using data loggers (Bonino et al. 2020, Pintanel et al. 2022), or simulated microclimate (Oyamaguchi et al. 2018). Understanding environmental tolerance across all life stages is crucial to identify vulnerability in for organisms with complex life cycles (Lowe et al. 2021). The differences in thermal tolerance between aquatic larval stages and mobile adult stages suggested that there is an evolutionary capacity to maximise performance to the environment for a given life stage. However, the potential to fully decouple detrimental environmental effects on fitness at one life stage to the next (i.e. carry-over effects) in amphibians needs to be explored at a comparative global scale.

## Acknowledgements

The authors would like to thank researchers C. Filho (Universidade Estadual de Santa Cruz) and R. Percinz (Ruth Percino) who provided data for this study.

## References

Anderson, R. O., S. Meiri, and D. G. Chapple. 2022. The biogeography of warming tolerance in lizards. Journal of Biogeography n/a:1274–1285.

Bates, A. E., and S. A. Morley. 2020. Interpreting empirical estimates of experimentally derived physiological and biological thermal limits in ectotherms. Canadian Journal of Zoology 98:237–244.

Bodensteiner, B. L., G. A. Agudelo-Cantero, A. Z. A. Arietta, A. R. Gunderson, M. M. Muñoz, J. M. Refsnider, and E. J. Gangloff. 2021. Thermal adaptation revisited: How conserved are thermal traits of reptiles and amphibians? Journal of Experimental Zoology Part A: Ecological and Integrative Physiology 335:173–194.

Bonino, M. F., F. B. Cruz, and M. G. Perotti. 2020. Does temperature at local scale explain thermal biology patterns of temperate tadpoles? Journal of Thermal Biology 94:102744.

Brattstrom, B. H. 1970. Thermal acclimation in Australian amphibians. Comparative Biochemistry and Physiology 35:69–103.

Bujan, J., K. A. Roeder, K. de Beurs, M. D. Weiser, and M. Kaspari. 2020. Thermal diversity of North American ant communities: Cold tolerance but not heat tolerance tracks ecosystem temperature. Global Ecology and Biogeography 29:1486–1494.

Burton, T., H.-K. Lakka, and S. Einum. 2020. Acclimation capacity and rate change through life in the zooplankton Daphnia. Proceedings of the Royal Society B: Biological Sciences 287:20200189.

Carilo Filho, L. M., B. T. de Carvalho, B. K. A. Azevedo, L.M. Gutiérrez-Pesquera, C.V. Mira-Mendes, M. Solé, and V. G. D. Orrico. 2021. Natural history predicts patterns of thermal vulnerability in amphibians from the Atlantic Rainforest of Brazil. Ecology and Evolution 11:16462–16472.

Casterlin, M. E., and W. W. Reynolds. 1978. Behavioural thermoregulation in Rana pipiens tadpoles. Journal of Thermal Biology 3:143–145.

Cheng, C.-T., M.-F. Chuang, T. Haramura, C.-B. Cheng, Y. I. Kim, A. Borzée, C.-S. Wu, Y.-H. Chen, Y. Jang, N. C. Wu, and Y.-C. Kam. 2022. Open habitats increase vulnerability of amphibian tadpoles to climate warming across latitude. Global Ecology and Biogeography n/a.

Chown, S. L., and K. J. Gaston. 2016. Macrophysiology – progress and prospects. Functional Ecology 30:330–344.

Collin, R., A. P. Rebolledo, E. Smith, and K. Y. K. Chan. 2021. Thermal tolerance of early development predicts the realized thermal niche in marine ectotherms. Functional Ecology 35:1679–1692.

Cupp, P. V. 1980. Thermal Tolerance of Five Salientian Amphibians during Development and Metamorphosis. Herpetologica 36:234–244.

Dahlke, F. T., S. Wohlrab, M. Butzin, and H.-O. Pörtner. 2020. Thermal bottlenecks in the life cycle define climate vulnerability of fish. Science 369:65–70.

Deutsch, C. A., J. J. Tewksbury, R. B. Huey, K. S. Sheldon, C. K. Ghalambor, D. C. Haak, and P. R. Martin. 2008. Impacts of climate warming on terrestrial ectotherms across latitude. Proceedings of the National Academy of Sciences of the United States of America 105:6668–6672.

Duarte, H., M. Tejedo, M. Katzenberger, F. Marangoni, D. Baldo, J. F. Beltrán, D.A. Martí, A. Richter-Boix, and A. Gonzalez-Voyer. 2012. Can amphibians take the heat? Vulnerability to climate warming in subtropical and temperate larval amphibian communities. Global Change Biology 18:412–421.

Enriquez-Urzelai, U., M. Sacco, A. S. Palacio, P. Pintanel, M. Tejedo, and A. G. Nicieza. 2019. Ontogenetic reduction in thermal tolerance is not alleviated by earlier developmental acclimation in Rana temporaria. Oecologia 189:385–394.

Enriquez-Urzelai, U., R. Tingley, M. R. Kearney, M. Sacco, A. S. Palacio, M. Tejedo, and A. G. Nicieza. 2020. The roles of acclimation and behaviour in buffering climate change impacts along elevational gradients. Journal of Animal Ecology 89:1722–1734.

Fick, S. E., and R. J. Hijmans. 2017. WorldClim 2: new 1-km spatial resolution climate surfaces for global land areas. International Journal of Climatology 37:4302–4315.

Forster, J., and A. G. Hirst. 2012. The temperature-size rule emerges from ontogenetic differences between growth and development rates. Functional Ecology 26:483–492.

Gahm, K., A. Z. A. Arietta, and D. K. Skelly. 2021. Temperature-mediated trade-off between development and performance in larval wood frogs (Rana sylvatica). Journal of Experimental Zoology Part A: Ecological and Integrative Physiology 335:146–157.

Garcia, R. A., J. L. Allen, and S. Clusella-Trullas. 2019. Rethinking the scale and formulation of indices assessing organism vulnerability to warmer habitats. Ecography 42:1024–1036.

González-del-Pliego, P., B. R. Scheffers, R. P. Freckleton, E. W. Basham, M. B. Araújo, A. R. Acosta-Galvis, C. A. Medina Uribe, T. Haugaasen, and D. P. Edwards. 2020. Thermal tolerance and the importance of microhabitats for Andean frogs in the context of land use and climate change. Journal of Animal Ecology 89:2451–2460.

Greer, A. 1980. Critical Thermal Maximum Temperatures in Australian Scincid Lizards: Their Ecological and Evolutionary Significance. Australian Journal of Zoology 28:91–102.

Gutiérrez-Pesquera, L. M., M. Tejedo, M.Á. Olalla-Tárraga, H. Duarte, A. Nicieza, and M. Solé. 2016. Testing the climate variability hypothesis in thermal tolerance limits of tropical and temperate tadpoles. Journal of Biogeography 43:1166–1178.

Haddad, C. F. B., and C. P. A. Prado. 2005. Reproductive Modes in Frogs and Their Unexpected Diversity in the Atlantic Forest of Brazil. Bioscience 55:207–217.

Hutchison, V. H. 1961. Critical Thermal Maxima in Salamanders. Physiological Zoology 34:92–125.

IUCN. 2022. The IUCN Red List of Threatened Species.

Jetz, W., and R. A. Pyron. 2018. The interplay of past diversification and evolutionary isolation with present imperilment across the amphibian tree of life. Nature Ecology & Evolution 2:850–858.

Katzenberger, M., J. Hammond, M. Tejedo, and R. Relyea. 2018. Source of environmental data and warming tolerance estimation in six species of North American larval anurans. Journal of Thermal Biology 76:171–178.

Kearney, M. R. 2013. Activity restriction and the mechanistic basis for extinctions under climate warming. Ecology Letters 16:1470–1479.

Kearney, M. R., and W. P. Porter. 2017. NicheMapR - an R package for biophysical modelling: the microclimate model. Ecography 40:664–674.

Kearney, M. R., and W. P. Porter. 2019. NicheMapR – an R package for biophysical modelling: the ectotherm and Dynamic Energy Budget models. Ecography 43:85–96.

Kingsolver, J. G., and L. B. Buckley. 2020. Ontogenetic variation in thermal sensitivity shapes insect ecological responses to climate change. Current Opinion in Insect Science 41:17–24.

Klockmann, M., F. Günter, and K. Fischer. 2017. Heat resistance throughout ontogeny: body size constrains thermal tolerance. Global Change Biology 23:686–696.

Leahy, L., B. R. Scheffers, S. E. Williams, and A. N. Andersen. 2022. Arboreality drives heat tolerance while elevation drives cold tolerance in tropical rainforest ants. Ecology 103:e03549.

Lenoir, J., T. Hattab, and G. Pierre. 2017. Climatic microrefugia under anthropogenic climate change: implications for species redistribution. Ecography 40:253–266.

Levy, O., L. B. Buckley, T. H. Keitt, C. D. Smith, K. O. Boateng, D. S. Kumar, and M. J. Angilletta, Jr. 2015. Resolving the life cycle alters expected impacts of climate change. Proceedings: Biological Sciences 282:20150837.

Lowe, W. H., T. E. Martin, D. K. Skelly, and H. A. Woods. 2021. Metamorphosis in an Era of Increasing Climate Variability. Trends in Ecology & Evolution 36:360–375.

Michelangeli, M., C. T. Goulet, H. S. Kang, B. B. M. Wong, and D. G. Chapple. 2018. Integrating thermal physiology within a syndrome: Locomotion, personality and habitat selection in an ectotherm. Functional Ecology 32:970–981.

Moen, D. S., E. Cabrera-Guzman, I. W. Caviedes-Solis, E. Gonzalez-Bernal, and A. R. Hanna. 2022. Phylogenetic analysis of adaptation in comparative physiology and biomechanics: overview and a case study of thermal physiology in treefrogs. Journal of Experimental Biology 225:jeb243292.

Moen, D. S., and J. J. Wiens. 2017. Microhabitat and Climatic Niche Change Explain Patterns of Diversification among Frog Families. The American Naturalist 190:29–44.

Morley, S. A., L. S. Peck, J. Sunday, S. Heiser, and A. E. Bates. 2018. Acclimation potential of global ectothermic species, collated from literature, 1960 to 2015. Natural Environment Research Council, Cambridge, UK, Polar Data Centre.

Moyano, M., C. Candebat, Y. Ruhbaum, S. Álvarez-Fernández, G. Claireaux, J.-L. Zambonino-Infante, and M. A. Peck. 2017. Effects of warming rate, acclimation temperature and ontogeny on the critical thermal maximum of temperate marine fish larvae. PLOS ONE 12:e0179928.

Noble, D. W. A., P. Pottier, M. Lagisz, S. Burke, S. M. Drobniak, R. E. O’Dea, and S. Nakagawa. 2022. Meta-analytic approaches and effect sizes to account for ‘nuisance heterogeneity’ in comparative physiology. Journal of Experimental Biology 225:jeb243225.

Oyamaguchi, H. M., P. Vo, K. Grewal, R. Do, E. Erwin, N. Jeong, K. Tse, C. Chen, M. Miyake, A. Lin, and M. Gridi-Papp. 2018. Thermal sensitivity of a Neotropical amphibian (Engystomops pustulosus) and its vulnerability to climate change. Biotropica 50:326–337.

Pandori, L. L. M., and C. J. B. Sorte. 2019. The weakest link: sensitivity to climate extremes across life stages of marine invertebrates. Oikos 128:621–629.

Pie, M. R., L. L. F. Campos, A. L. S. Meyer, and A. Duran. 2017. The evolution of climatic niches in squamate reptiles. Proceedings: Biological Sciences 284.

Pincebourde, S., and J. Casas. 2015. Warming tolerance across insect ontogeny: influence of joint shifts in microclimates and thermal limits. Ecology 96:986–997.

Pincebourde, S., and H. A. Woods. 2020. There is plenty of room at the bottom: microclimates drive insect vulnerability to climate change. Current Opinion in Insect Science 41:63–70.

Pinsky, M. L., A. M. Eikeset, D. J. McCauley, J. L. Payne, and J. M. Sunday. 2019. Greater vulnerability to warming of marine versus terrestrial ectotherms. Nature 569:108–111.

Pintanel, P., M. Tejedo, A. Merino-Viteri, F. Almeida-Reinoso, S. Salinas-Ivanenko, A. C. Lopez-Rosero, G. A. Llorente, and L. M. Gutierrez-Pesquera. 2022. Elevational and local climate variability predicts thermal breadth of mountain tropical tadpoles. Ecography 2022:e05906.

Pintanel, P., M. Tejedo, S. R. Ron, G. A. Llorente, and A. Merino-Viteri. 2019. Elevational and microclimatic drivers of thermal tolerance in Andean Pristimantis frogs. Journal of Biogeography 46:1664–1675.

Rohr, J. R., D. J. Civitello, J. M. Cohen, E. A. Roznik, B. Sinervo, and A. I. Dell. 2018. The complex drivers of thermal acclimation and breadth in ectotherms. Ecology Letters 21:1425–1439.

Rubalcaba, J. G., and M.Á. Olalla-Tárraga. 2020. The biogeography of thermal risk for terrestrial ectotherms: Scaling of thermal tolerance with body size and latitude. Journal of Animal Ecology n/a:1277–1285.

Ruthsatz, K., K. H. Dausmann, M. A. Peck, and J. Glos. 2022. Thermal tolerance and acclimation capacity in the European common frog (Rana temporaria) change throughout ontogeny. J Exp Zool A Ecol Integr Physiol 337:477–490.

Sinclair, B. J., K. E. Marshall, M. A. Sewell, D. L. Levesque, C. S. Willett, S. Slotsbo, Y. Dong, C. D. Harley, D. J. Marshall, B. S. Helmuth, and R. B. Huey. 2016. Can we predict ectotherm responses to climate change using thermal performance curves and body temperatures? Ecology Letters 19:1372–1385.

Sunday, J. M., A. E. Bates, M. R. Kearney, R. K. Colwell, N. K. Dulvy, J. T. Longino, and R. B. Huey. 2014. Thermal-safety margins and the necessity of thermoregulatory behavior across latitude and elevation. Proceedings of the National Academy of Sciences of the United States of America 111:5610–5615.

Tracy, C. R., K. A. Christian, and C. R. Tracy. 2010. Not just small, wet, and cold: effects of body size and skin resistance on thermoregulation and arboreality of frogs. Ecology 91:1477–1484.

Truebano, M., P. Fenner, O. Tills, S. D. Rundle, and E. L. Rezende. 2018. Thermal strategies vary with life history stage. Journal of Experimental Biology 221:jeb171629.

Vidal-García, M., and J. S. Keogh. 2015. Convergent evolution across the Australian continent: ecotype diversification drives morphological convergence in two distantly related clades of Australian frogs. Journal of Evolutionary Biology 28:2136–2151.

Wilson, E. S., C. E. Murphy, J. P. Rinehart, G. Yocum, and J. H. Bowsher. 2020. Microclimate Temperatures Impact Nesting Preference in Megachile rotundata (Hymenoptera: Megachilidae). Environmental Entomology.

Wu, N. C., R. P. Bovo, U. Enriquez-Urzelai, S. Clusella-Trullas, M. R. Kearney, C. A. Navas, and J. D. Kong. 2024. Global exposure risk of frogs to increasing environmental dryness. Nature Climate Change 14.

